# Leafhopper transmits soybean stay-green associated virus to legume plants

**DOI:** 10.1101/2023.01.05.522956

**Authors:** Jinlong Yin, Zhuangzhuang Hu, Shuqi Xu, Xin Hong, Yanglin Qiu, Xinge Cheng, Liqun Wang, Weiliang Shen, Haijian Zhi, Kai Li, Kai Xu

**Author notes:** Correspondence (Kai Xu); (Kai Li).

## Abstract

A novel geminivirus, soybean stay-green associated virus (SoSGV), was previously shown to cause soybean delayed senescence and is associated with the incidences of soybean stay-green syndrome. The transmission methods of SoSGV were not yet understood. We captured insects belonging to 24 distinct species in a soybean field with the SoSGV outbreak and detected the presence of SoSGV only in leafhoppers and bean bugs (*Riptortus pedestris*). Caged feeding experiments using captured leafhoppers and bean bugs from soybean fields showed that leafhoppers, but not bean bugs, are vectors transmitting SoSGV. The common brown leafhopper (*Orosius orientalis*) is identified as the dominant leafhopper species and can establish colonies feeding on soybean plants in experimental conditions. An investigation of SoSGV defective DNA revealed that soybean genomic DNA could be inserted into the SoSGV genome, while sequences from wild soybean, red bean, and cowpea were also identified. We further showed that the common brown leafhopper could transmit SoSGV to wild soybean and red bean plants, emphasizing the vector role of the leafhopper in the transmission of SoSGV in the field.

## Introduction

Soybean Stay-Green Syndrome (SGS), commonly called ‘Zhengqing’ in Chinese, is a soybean disorder involving delayed senescence of the whole or parts of the plant. In China, SGS occurred during the 1980s in the Huang-Huai-Hai Plain and become a significant issue in soybean production by causing severe yield loss [1]. In the United States of America, soybean delayed senescence, also known as the green stem syndrome, was recorded as early as 1950 and is still a problem for soybean growers [2]. Similar maturity disorder of soybean has also been reported in soybean-growing countries like Brazil, Argentine, and Japan [3–5].

Despite delayed senescence being the key feature of SGS, the accompanying symptoms vary depending on the year and location. In China, a briefing in 2002 described SGS plants in Luohe (Henan province) as reduced pod number, continued vegetative growth in the late season, and swollen seeds [6]. The reduced pod number caused by the falling of blooms/pods was attributed as a typical symptom of several latter occurred SGS incidences in Henan province [7–9]. Starting in the 2010s, the poor seed quality caused by a high percentage of abnormal pod-filling during the R7-R8 stage became a major indicator of SGS [10–12], while the decrease in pod number seems not evident in some studies [12, 13]. In addition to the pod number and quality, the SGS plants in the Huang-Huai-Hai Plain region can have either normal height and leaf shape or sometimes be stunted with curling and crinkling leaf shapes [12]. In one given SGS plant, the delayed maturity can occur in the whole plant or only occur in a part of the plant [10]. In the USA, different symptoms were observed for delayed senescence of soybean [2]. One major type, often described by terms like green plant malady, green stem malady, green bean syndrome, or green stem syndrome, is characterized by reduced pod number, poor seed quality, immature plant parts, and yield loss. The other type, called green stem disorder, is associated with green stem phenotype and normal seed quality and maturity. In Brazil, at least two types of soybean maturity disorder were seen, distinguished by whether the leaves, stems, and pods are deformed [3].

Currently, the causal agent for delayed senescence is still not perspicuous. The pod removal experiments suggested that the imbalance of leaf-seed (source-sink) relationships causes delayed senescence or SGS [14, 15], but natural pod removal usually does not occur, and other factors were thought to trigger the source-sink imbalance [16]. The feeding of bean bug *Riptortus pedestris* (Fabricius) (*Hemiptera: Alydidae*) is shown to be one of the possible causes of SGS in the Huang-Huai-Hai region of China [12, 17], as the feeding of bean bugs on the R3 – R5 stage soybean plants in the field cage or the greenhouse can lead to seed damage and SGS symptoms [12, 17–19]. Similarly, the feeding of red-banded stink bug *Piezodorus guildinii* (Westwood) (*Hemiptera: Pentatomidae*) with R4 stage soybeans was also found to cause flat pods, green leaf retention, and yield loss based on a study performed in Texas, USA [20]. In Brazil, the leaf nematode *Aphelenchoides besseyi* causes the delayed senescence of soybean that is associated with viral disease-like leaf distortion and vein thickening symptoms [3]. Some studies linked RNA viruses, bean leaf beetle, fungicide spray, and light or water conditions with soybean delayed senescence [2, 4, 21–25]. However, no single reason seems to account for this global soybean growing issue, but particular factors may be the cause in a given area during specific years [16, 26].

Recently, a novel geminivirus, soybean stay-green associated virus (SoSGV) [13], has been identified and found to be strongly associated with the SGS incidences in the Huang-Huai-Hai region of China [13, 27–29]. Inoculation of this virus to soybean plants based on an infectious viral clone recapitulates the SGS symptoms, like increased flat pod number and reduced yield, while the total pod number keeps unchanged [13]. However, the transmission method of this virus is currently unknown. Given that insecticide spray can significantly reduce the SGS occurrence in the fields of the Huang-Huai-Hai region [1, 17, 30, 31], it is worth investigating whether insect vector can transmit SoSGV.

In this study, we captured 24 species of insects that infested the SoSGV-infected soybean plants. These insects were then subjected to total DNA extraction followed by the Polymerase Chain Reaction (PCR) to detect the presence of SoSGV. We found that SoSGV can only be detected in the leafhopper *Orosius sp. (Hemiptera: Cicadellidae*) and in the bean bug *R. pedestris*. Caged feeding experiments demonstrated that the leafhoppers, but not the bean bugs, transmit SoSGV to healthy soybean seedlings and other legume plants. Our results provided key knowledge of the SoSGV epidemic and will aid a better understanding of the SGS occurrence in the Huang-Huai-Hai region of China.

## Results

### New SoSGV isolates identified in soybean stay-green plants

During the autumn of 2022, we collected soybean samples that showed stay-green symptoms from major soybean-growing provinces/municipalities of China, including Anhui, Jiangsu, Shanxi, Shaanxi, and Beijing. Based on PCR detection using SoSGV-specific primer pairs, 31 out of 34 samples from Anhui, 10 out of 13 samples from Jiangsu, 3 out of 5 samples from Shanxi, and 27 out of 30 samples from Beijing tested positive for SoSGV (Supplementary Table 2), of which Jiangsu, Shanxi, and Beijing are previously unknown for the epidemic of SoSGV. We detected no SoSGV infection in SGS samples collected in Shaanxi Province. Combining previously reported locations, SoSGV seems to spread widely in the Huang-Huai-Hai region of China (Fig. 1A).

**Figure 1.**
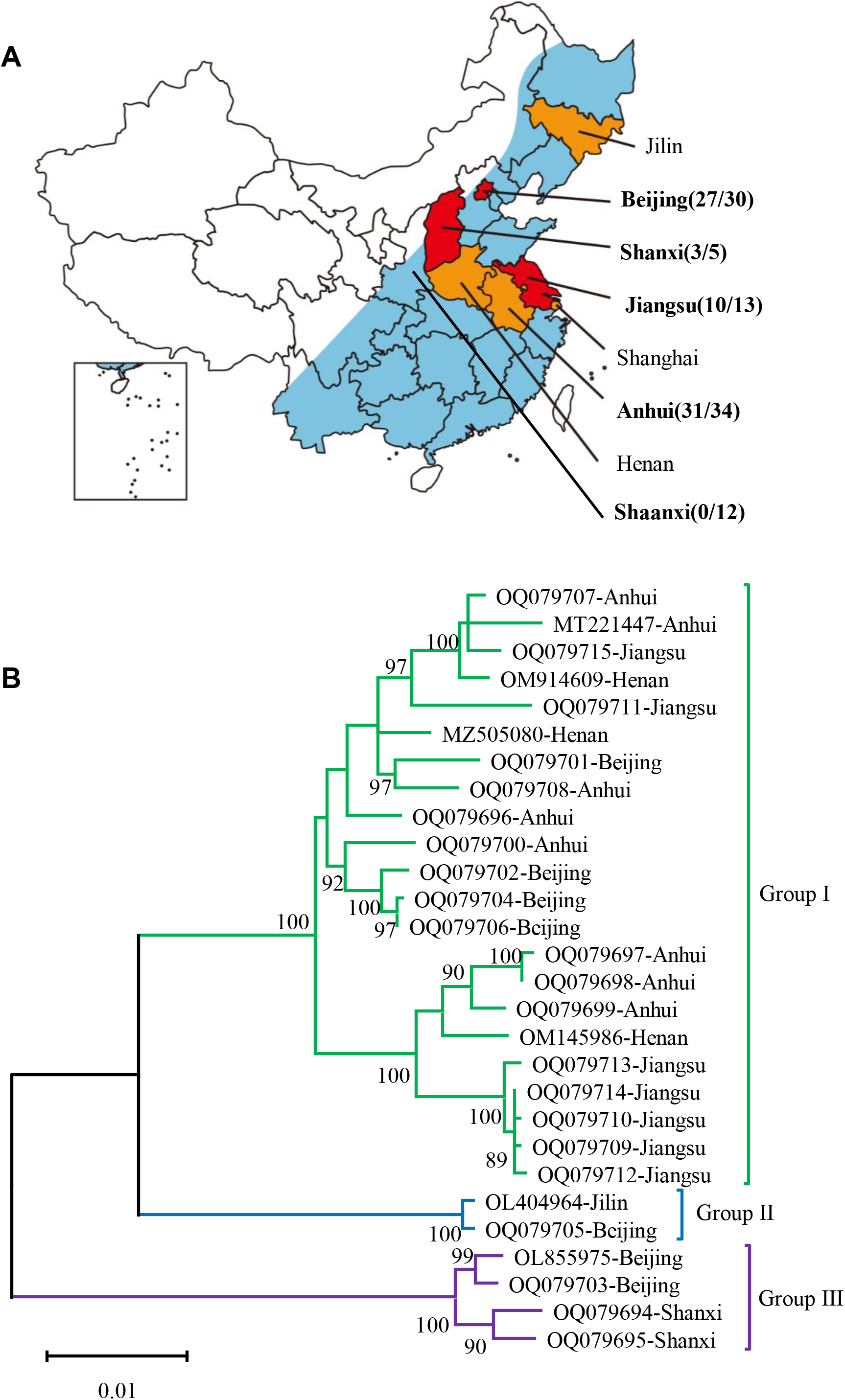
Distribution and phylogenetics of SoSGV in China. (A) Distribution of SoSGV identified in various locations in China. The blue-colored region in the map represents China’s main soybean production areas. The yellow-colored areas represent the provinces where SoSGV was identified in previous reports. The red-colored areas are the newly reported epidemic areas in this study. The locations where the samples were obtained in this study are written in bold font, followed by the number of SoSGV-positive samples over the total number of SGS samples collected (positive samples/total samples). (B) The unrooted phylogenetic tree was constructed based on the nucleotide sequence of the full-length genome of different SoSGV isolates. The tree was built with MEGA (version 11.0.11). The branch lengths represent the substitution rate per nucleotide site. Bootstrap values higher than 70 were given at the nodes. The accession numbers and the sampling locations are given for each isolate. Those 22 isolates with accession numbers OQ079694 - OQ079715 are sequenced and reported in this study.

Some SoSGV-positive samples were chosen for PCR amplification of the full-length SoSGV genome, followed by sanger sequencing. The obtained genome sequences showed more than 93% nucleotide identity with each other or with all six currently-available SoSGV sequences from the NCBI database (Supplementary Fig. 1). Phylogenetic analysis of the overall 28 SoSGV sequences revealed a total of 3 evolutionary groups. Group I mainly consists of the isolates from the Huang-Huai-Hai region, including Henan, Anhui, and Jiangsu. Group II is represented by isolates from Jilin and Beijing, while Group III contains Shanxi and Beijing isolates. The Huang-Huai-Hai Plain is a confined area physically separated from Shanxi province by Taihang Mountains, while the Yanshan Mountains and the Bohai Gulf also distance the Huang-Huai-Hai area from the Northeast soybean-growing provinces, including Jilin. Thus, except for isolates from Beijing that seem to have actively been transmitted from neighboring regions, the three groups contain viral isolates of distinct geographic locations.

### Screen for insects carrying SoSGV

The wild-spread distribution of SoSGV is likely caused by insect transmission of the virus. To find out which insects carry and transmit SoSGV, we captured overall 24 insect species in a soybean field in Suzhou, Anhui Province, where SoSGV-positive soybean samples were previously collected. These insect species include those from Lepidoptera, Hemiptera, or Orthoptera (Fig. 2A). The total DNA was extracted from these insects and subsequently subjected to PCR using two pairs of SoSGV-specific primers (Supplementary Table 1). Among those 24 insect species, the leafhopper (*Orosius sp.*) tested positive by using one of the primer pairs (Fig. 2B, PCR1), while the bean bug (*R. pedestris*) tested positive using both primer pairs (Fig. 2B, PCR 1 and PCR 2). The other insects tested negative for SoSGV.

**Figure 2.**
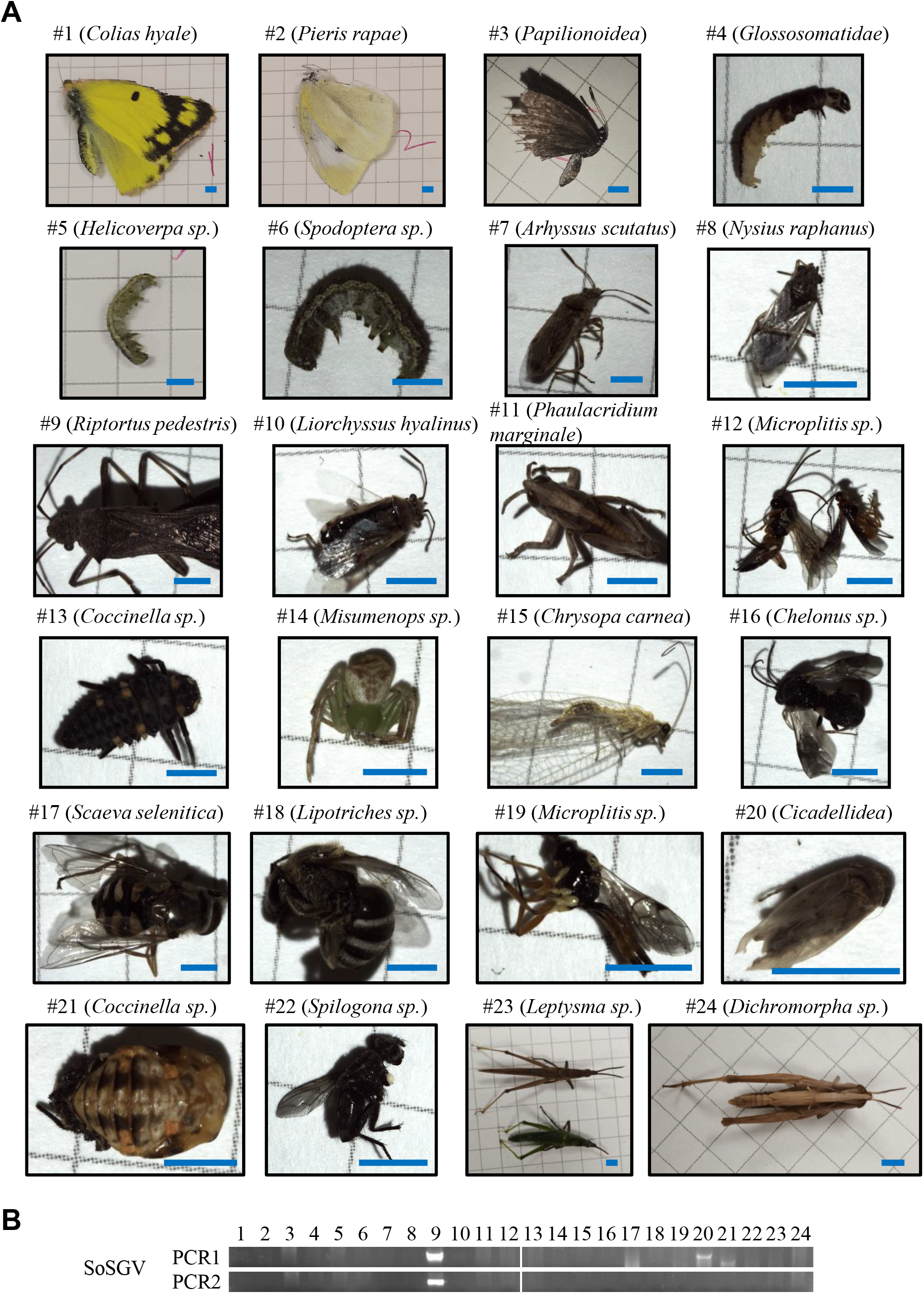
The screening for insects carrying SoSGV. (A) Images of the insects caught in a SoSGV-infected soybean field in Suzhou, Anhui Province. Scientific names at the rank of species, genus, or family were shown. Bar = 2.5 mm. (B) The detection of SoSGV by PCR from total DNA extracted from insects shown in panel A. Two primer pairs were used. Primers #2906 and #2907 were used for PCR 1. Primers #2904 and #2905 were used for PCR 2. The PCR products amplified from each insect were loaded to DNA gels in the order of insect species shown in panel A.

### Leafhoppers transmit SoSGV to healthy soybean plants

To test whether leafhopper can transmit SoSGV to soybean plants, we captured 50 leafhoppers from the field in Suzhou, Anhui Province, where the leafhoppers previously tested positive for SoSGV (Fig. 2). They were subsequently reared on 11 healthy V1 stage soybean seedlings in a meshed cage for 14 d. Leaf samples were then collected from individual plants at 12 days post-infestation (dpi), 15 dpi, and 34 dpi (Fig. 3A, 1st experiment). The collected leaf samples all tested positive for SoSGV, while the leaf samples from non-infested seedlings grown in a separate meshed cage tested negative (Fig. 3B). The leafhopper-infested soybean leaves are wrinkled and distorted (Fig. 3C). Among these 50 leafhoppers reared on soybean seedlings, only 13 leafhoppers survived after 14 d period of rearing. All surviving ones are common brown leafhopper *Orosius orientalis* (Matsumura) (Fig. 3D). It is likely that these common brown leafhoppers adapt to the diet of the soybean plants, while other leafhoppers can not survive solely on soybeans. Interestingly, SoSGV can still be detected in about half of the dead leafhoppers (Fig. 3E), suggesting that these leafhopper species may also carry and transmit SoSGV. In another repeat experiment, the 13 common brown leafhoppers were transferred to 8 healthy soybean seedlings in a new meshed cage (Fig. 3A). At 14 dpi, 6 out of 8 seedlings were infected by SoSGV, and at 21 dpi, all plants tested positive (Fig. 3F). These results demonstrate that leafhoppers, especially common brown leafhoppers, can transmit SoSGV to soybean plants.

**Figure 3.**
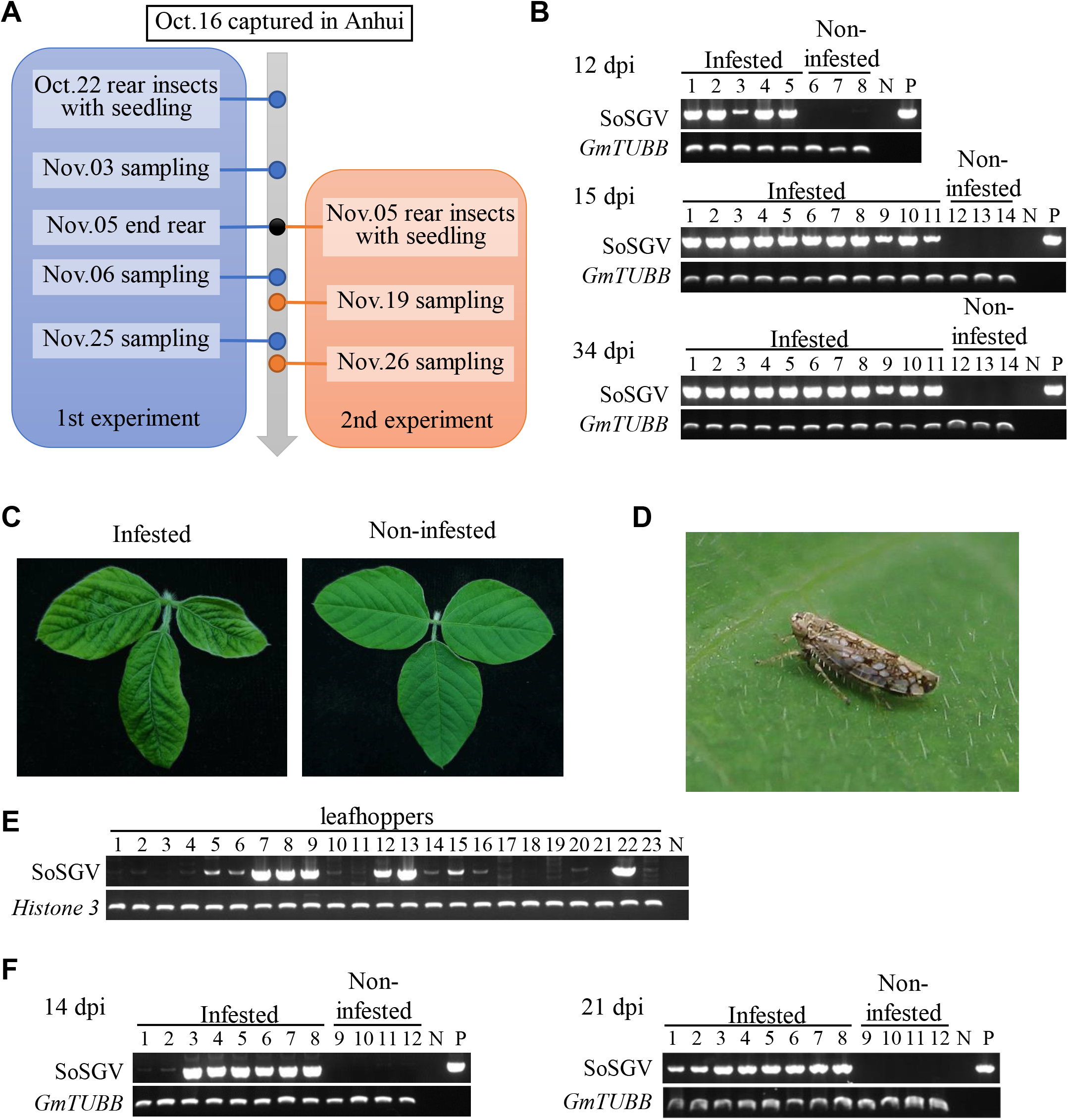
Leafhoppers captured in Anhui Province transmit SoSGV to soybean. (A) The time flow chart of two SoSGV transmission experiments. (B) PCR detecting SoSGV in soybean seedlings infested or non-infested by leafhoppers. SoSGV was amplified using primer pair #3133 and #3144. The gene coding for soybean β-tubulin (*GmTUBB*) was amplified via PCR as a loading control. Water and a plasmid containing SoSGV genome DNA were used as negative and positive controls, respectively. Different lanes represent different plants tested using PCR. (C) Leaf symptoms of soybean seedlings at 12 days post leafhopper infestation. Non-infested leaves were shown. (D) An adult common brown leafhopper (*O. orientalis*) feeding on a soybean leaf. (E) Detection of SoSGV in leafhoppers unable to survive the 14 d rearing on soybean seedlings. The gene coding for Histone 3 was used as a loading control. (F) PCR detecting SoSGV in soybean seedlings infested or non-infested by leafhoppers in the 2nd experiment. See further detail in panel B.

We also captured leafhoppers in a soybean field in Nanjing, Jiangsu Province, that had SGS plants previously tested positive for SoSGV infection (Fig. 1A). A total of 80 captured leafhoppers were reared on 8 soybean seedlings, all of which tested positive for SoSGV infection at 24 dpi (Fig. 4B). The experiment was repeated by rearing 9 common brown leafhoppers in a separate meshed cage with another 8 health seedlings. At 15 dpi, these seedlings are all infected by SoSGV despite the fact that 3 seedlings accumulated less SoSGV (Fig. 4C). The leafhopper-infested plants distorted in leaf shape (Fig. 4D).

**Figure 4.**
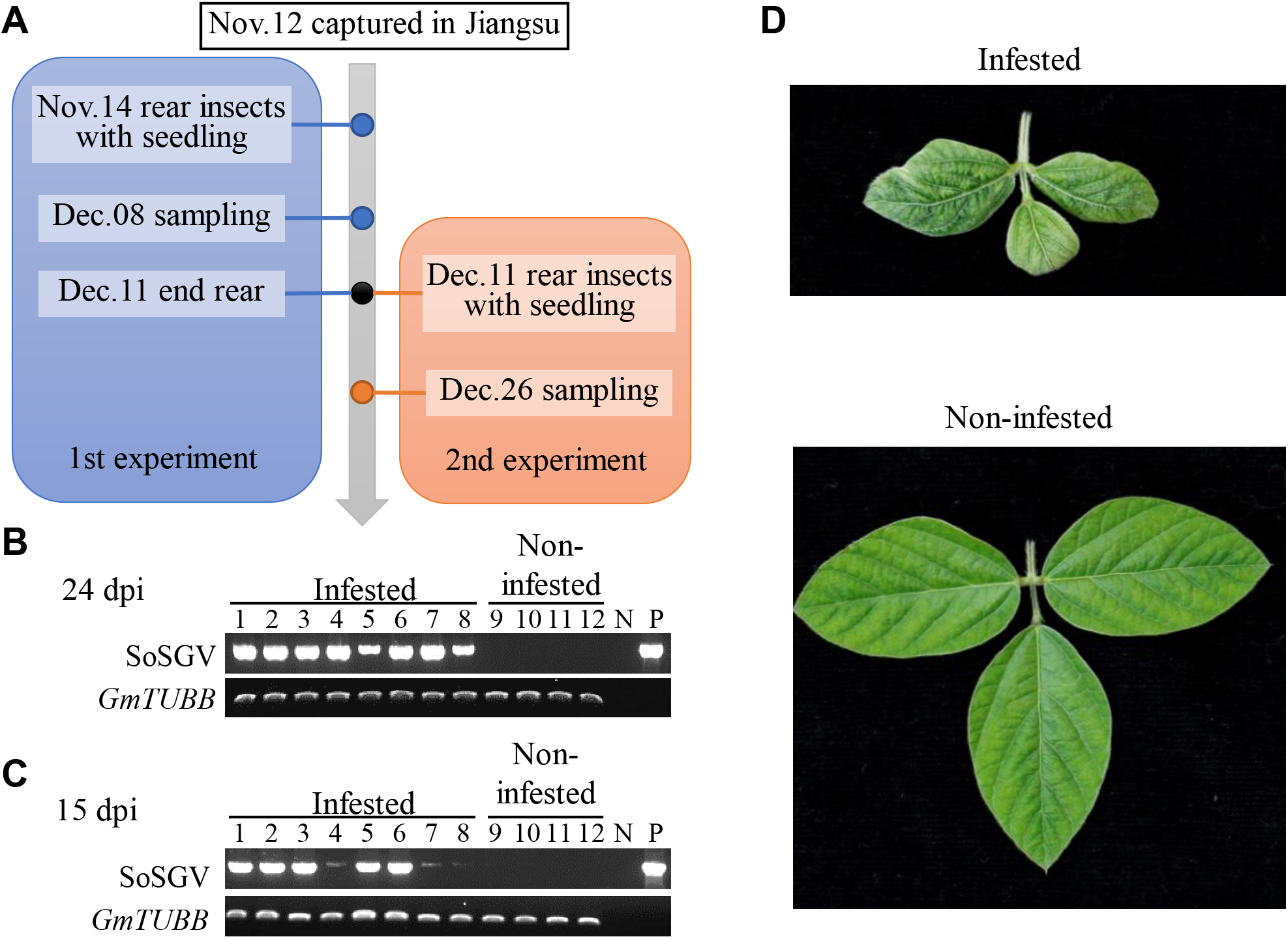
Leafhoppers captured in Jiangsu Province transmit SoSGV to soybean. (A) The time flow chart of two SoSGV transmission experiments. (B-C) PCR assay detecting SoSGV in soybean seedlings infested or non-infested by leafhoppers in two experiments depicted in panel A. Primers #3133 and #3144 were used to amplify SoSGV. The gene coding for soybean β-tubulin (*GmTUBB*) was amplified via PCR as a loading control. Water and a plasmid containing SoSGV genome DNA were used as negative and positive controls, respectively. Different lanes represent different plants tested using PCR. (D) Leaf symptoms of soybean seedlings at 24 days post leafhopper infestation. Non-infested leaves were shown.

### Bean bug *R. pedestris* does not transmit SoSGV

The bean bug *R. pedestris* is previously shown to induce SGS by feeding damage at the early pod stage [12, 17, 18]. The SGS caused by *R. pedestris* was shown to be independent of any virus infection [12]. We captured the *R. pedestris* adults from Suzhou, Anhui Province, and kept them in a meshed cage with soybean seedlings as a food source (Fig. 5A). In the first experiment, 25 *R. pedestris* adults were reared on 11 healthy V1 stage soybean seedlings for 13 days (Fig. 5B). These plants were continuously tested for the SoSGV infection at 12dpi, 15 dpi, 20 dpi, and 24 dpi. PCR and qPCR assay showed that some samples were weak positive at 12 dpi or 15 dpi, but none of the 11 plants were successfully infected by SoSGV at 24 dpi (Fig. 5C and 5D). The *R. pedestris* excretes a brownish and smelly liquid that usually stains the leaves. The weak band in the PCR assay might be due to the contamination of the insect excreta that contains the undigested SoSGV taken up in the field. In agreement with this observation, we found that although the *R. pedestris* can be detected positive for SoSGV after 9 days, the virus level significantly decreased after 35 days of feeding on seedlings or dry seeds (Fig. 5E). In the second experiment, 14 *R. pedestris* adults kept in the greenhouse for about half a month were transferred to a new meshed cage with 8 healthy soybean seedlings. The infested plants tested negative for SoSGV-infection after 14 or 21 days (Fig. 5F). Above results demonstrate that the bean bug *R. pedestris* can not transfer SoSGV in the experimental condition.

**Figure 5.**
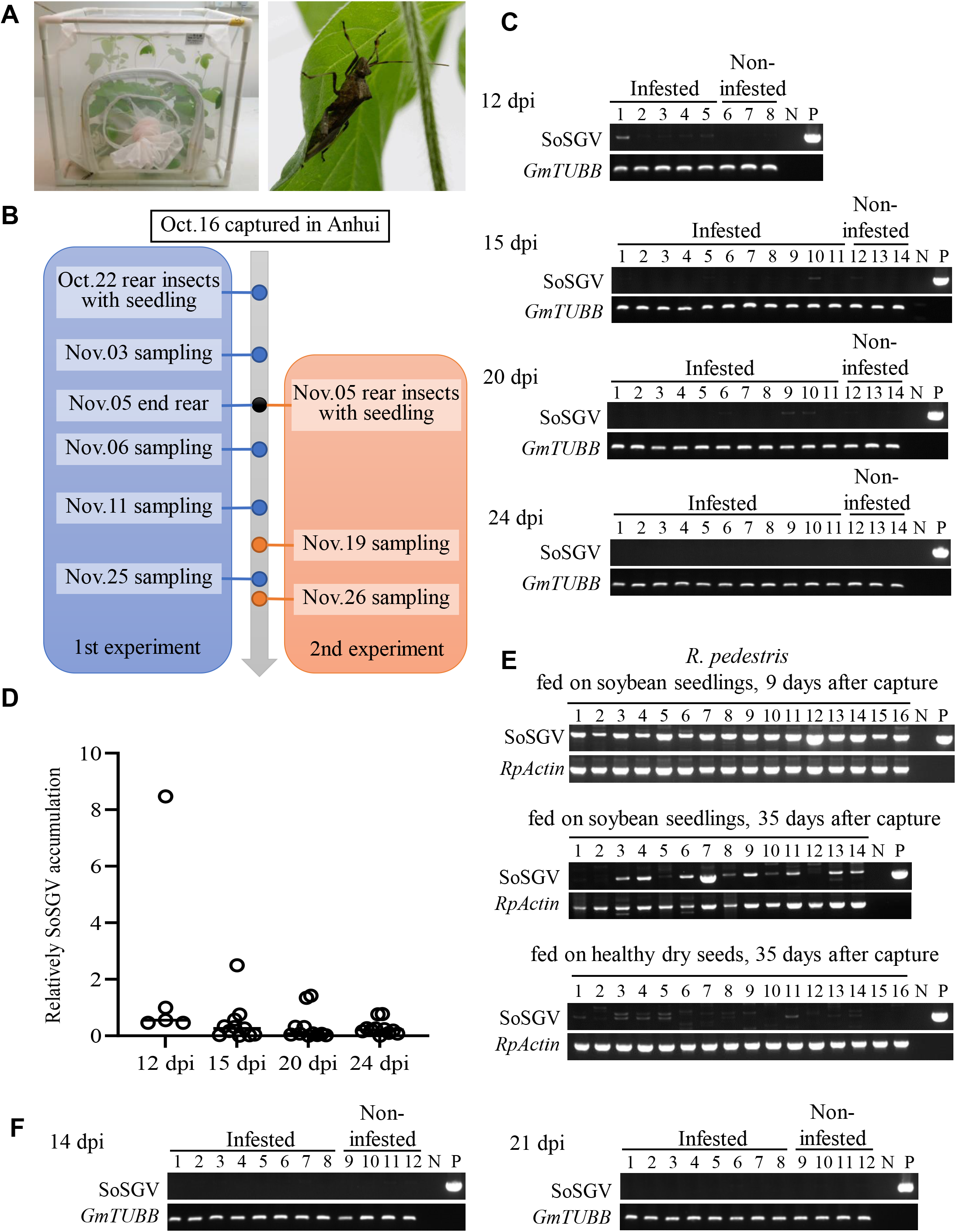
Bean bug (*Riptortus pedestris*) can not transmit SoSGV. (A) The experimental setup of SoSGV transmission assay. The insects were reared on soybean seedlings in a nylon cage. No other food sources were supplied. (B) The time flow chart of two SoSGV transmission experiments. (C) PCR detecting SoSGV in soybean seedlings infested or non-infested by *R. pedestris*. SoSGV was amplified using primer pair #3133 and #3144. The gene coding for soybean β-tubulin (*GmTUBB*) was amplified via PCR as a loading control. Water and a plasmid containing SoSGV genome DNA were used as negative and positive controls, respectively. Different lanes represent different plants tested using PCR. (D) quantitative PCR analysis of the same samples in panel C. (E) Detection of SoSGV in *R. pedestris* via PCR. The *R. pedestris* adults were either reared on soybean seedlings or on dry soybean seeds at the indicated time. The gene coding for *R. pedestris* actin (*RpActin*) was used as a loading control. (F) PCR detecting SoSGV in soybean seedlings infested or non-infested by *R. pedestris* in the 2nd experiment. See further detail in panel C.

### Defective SoSGV contains genomic DNA fragments from legume plants

When the full-length viral genome (~ 2700 bp) was amplified via PCR from SoSGV-infected soybean samples using back-to-back primers (#2914 and #2915), a thick band of ~1000 bp was also amplified (Fig. 6A and 6B). Sanger sequencing of the gel-isolated PCR products suggested that the ~1000 bp amplicon represents the defective DNA molecules of SoSGV (Supplementary Table 4). The defective DNA occasionally contains non-SoSGV sequences located between the C1 coding region and the big interval, while lacking the C2, C3, V1, V2, and a small part of the C1 coding sequences (Fig. 6C). To investigate the origin of the non-SoSGV sequences, the PCR products containing this region was amplified from SoSGV-infected soybean samples collected from Anhui, Beijing, and Shanxi (with primers #3059 and #3058) and subjected to high throughput sequencing. A total of 170 thousand pairs of reads from each end of the PCR amplicon were sequenced. Each paired read was assembled and processed to remove those of SoSGV-origin, non-specifically amplified, or repeated reads. The generated 242 unique reads containing non-SoSGV sequences were searched against the non-virus nr database of NCBI by BLAST. Among them, 69 are sequences of soybean (*Glycine max*) genomic DNA, 2 hit wild soybean (*Glycine soja*), 2 hit cowpea (*Vigna unguiculata*), and 1 hits red bean (*Vigna angularis*) (Fig. 6D and Supplementary Table 5). As shown in Fig. 6E, the underlined sequence is 83 bp in length and identical to an 83 bp region in the reported *G. soja* sequence (Genbank accession # XR_003658818). The presence of plant sequences in SoSGV-infected soybean samples could also be verified by PCR (Supplementary Figures 2, 3, and 4).

**Figure 6.**
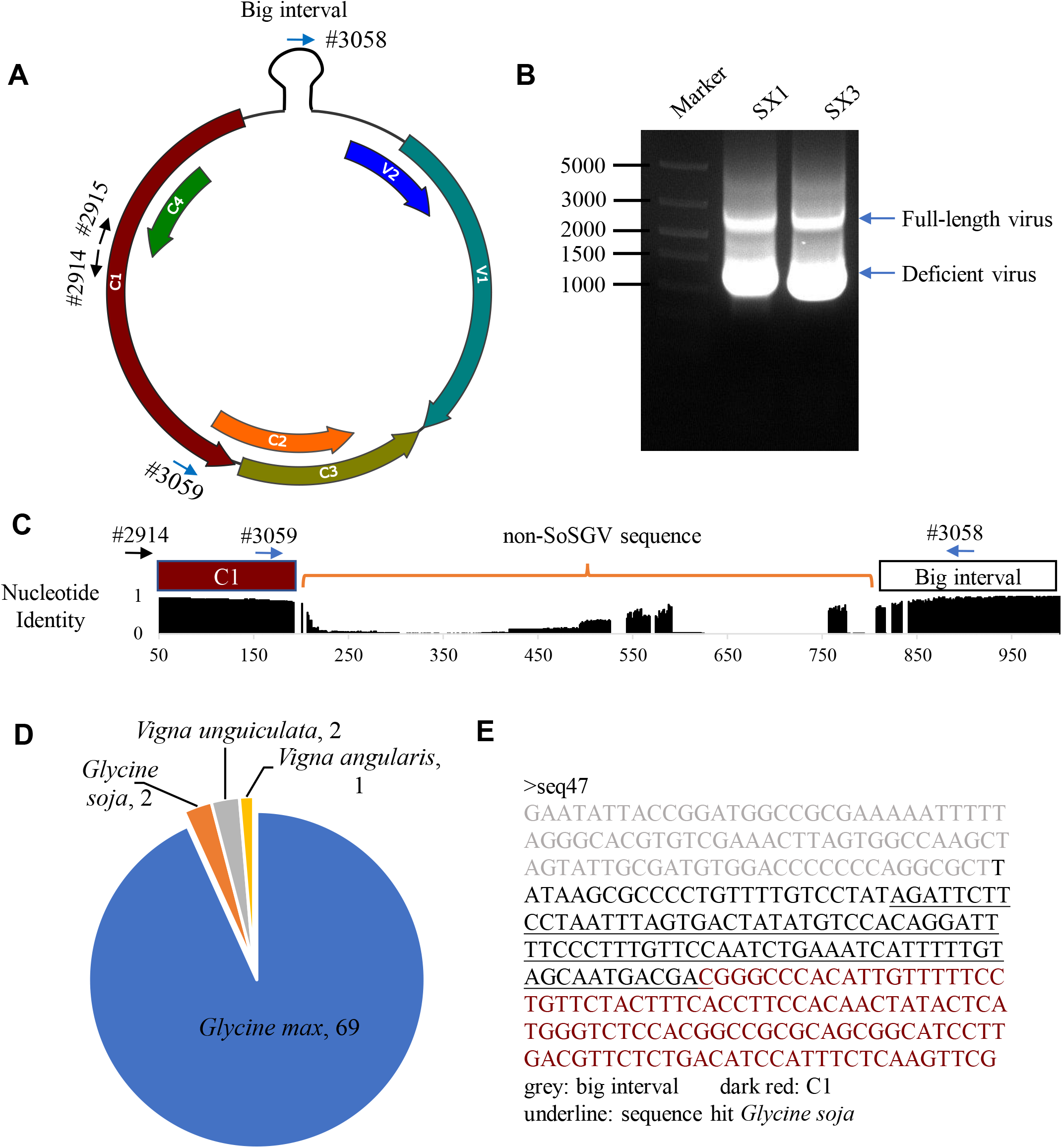
SoSGV defective DNAs contain host sequences. (A) Schematic representation of the SoSGV genome structure. (B) PCR products of amplified SoSGV genomic DNAs separated in agarose gel electrophoresis. The PCR is performed using back-to-back primers (#2914 and #2915), as shown in panel A. The PCR products of two samples from Shanxi province are shown. (C) An identity histogram of the aligned 75 defective DNA obtained from sanger sequencing of the cloned gel-isolated PCR products. The non-SoSGV sequences were found between the C1 and the big interval coding sequences. Primers #3059 and #3058 were used to amplify the non-SoSGV sequence-containing region for high-throughput sequencing. (D) A pie chart showing the numbers of unique sequences matching sequences of different plant species based on high-throughput sequencing. Stay-green soybean samples from Anhui, Shanxi, and Beijing were subjected to high-throughput sequencing of the non-SoSGV sequence. (E) A DNA fragment from wild soybean inserted in the SoSGV defective DNA clone seq47 is shown as an example.

The insertion of specific plant DNA sequences into the SoSGV might happen during the viral genome replication. Our finding indicates that the wild soybean, cowpea, or red bean could host the replication of SoSGV in the field, where the insect vectors might mediate the transmission of SoSGV among them.

### Leafhopper transmits SoSGV to wild soybean and red bean

To test whether SoSGV can infect wild soybean, red bean, or cowpea and whether leafhoppers can mediate the viral transmission to these plants, we transferred ~50 common brown leafhopper nymphs born and fed on SoSGV-infected soybean plants (Fig. 7A) onto wild soybean, red bean, or cowpea seedlings in a meshed cage. The wild soybean *G. soja* var. ‘Yong-40’ was more susceptible to SoSGV than ‘Yong-10’ and ‘Yong-27’ since all the ‘Yong-40’ seedlings were infected at 14 dpi, while only 1 ‘Yong-27’ seedling was infected. At 27 dpi, 3 out of 6 seedlings of ‘Yong-10’ and 4 out of 6 seedlings of ‘Yong-27’ were infected (Fig. 7B). The leafhopper-infested and SoSGV-infected wild soybean plants showed deformed leaves (Fig. 7C). The leafhopper infested red bean (*V. unguiculata*) were all infected at 15 dpi (Fig. 7D) and showed stunt growth (Fig. 7E). In contrast, the cowpea *V. unguiculata* cv. ‘Te xuan zhang tang wang’ we used in this study was resistant to SoSGV, as none of the seedlings were infected even at 27 dpi. These results demonstrate that the leafhopper could mediate the transmission of SoSGV to legume plants.

**Figure 7.**
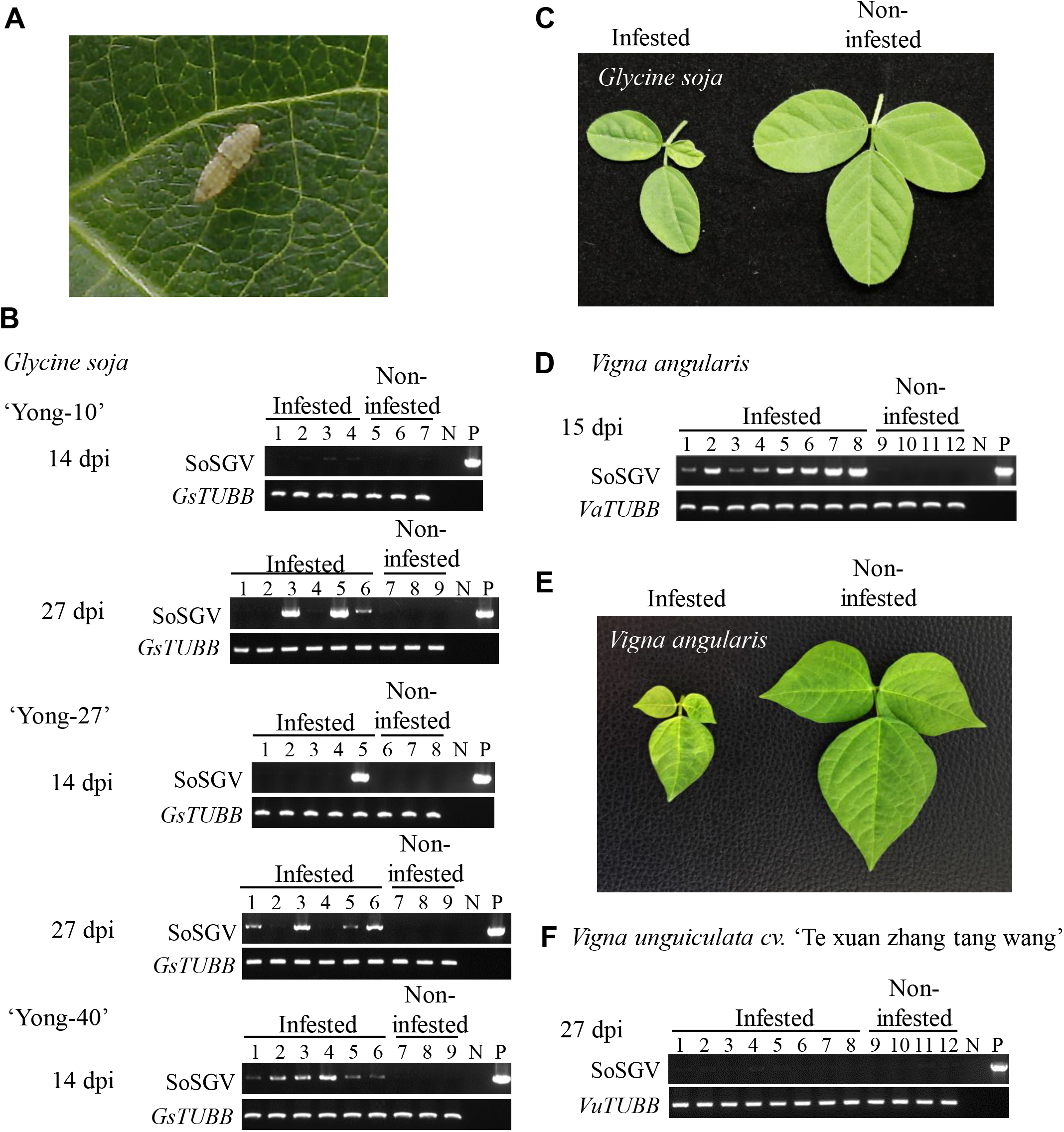
Common brown leafhopper (*O. orientalis*) transmits SoSGV to other legume plants. (A) Image of an *O. orientalis* nymph feeding on a soybean leaf. (B) PCR detecting SoSGV in different varieties of wild soybean *Glycine soja* infested or non-infested by nymphs of the common brown leafhoppers. SoSGV was amplified using primer pair #3133 and #3144. The gene coding for β-tubulin (*GsTUBB*) was amplified via PCR as a loading control. Water and a plasmid containing SoSGV genome DNA were used as negative and positive controls, respectively. Different lanes represent different plants tested using PCR. (C) Leaf symptoms of wild soybean variety ‘Yong-40’ at 14 days post-*O. orientalis* infestation. (D) PCR detection of SoSGV in a *Vigna angularis* landrace. See further details in panel B. (E) Leaf symptoms of *Vigna angularis* at 15 days post-*O. orientalis* infestation. (F) PCR detection of SoSGV in cowpea (*Vigna unguiculata*) cv. ‘Te xuan zhang tang wang’. See further details in panel B.

## Discussion

The Soybean SGS in China has caught the attention of growers and researchers for decades [1, 6–8, 10–12, 14, 17, 30, 31]. The geminivirus SoSGV was previously shown to be highly correlated to the incidences of SGS in the field and could cause SGS symptoms when inoculated to the soybean plants in the greenhouse [13, 28]. We found that the SGS is associated with SoSGV, with exceptions from the SGS samples collected in Shaanxi (Fig. 1A and Supplementary Table 2). It will be interesting to determine whether the bean bug or other reasons cause SGS in the soybean fields in Shaanxi Province. Meanwhile, most of the SGS samples collected from Anhui, Jiangsu, Shanxi, and Beijing were infected by SoSGV, but a few of them still tested negative (Fig. 1A and Supplementary Table 2). Although we can not exclude other possibilities, including that the samples taken for PCR detection were from the healthy parts of SoSGV-infected plants, the *R. pedestris* infestation should not be excluded as a cause for these SGS incidences with high SoSGV-infection rates. In fact, *R. pedestris* infestation were also observed in the soybean fields where we collected the SGS samples in Anhui, Shanxi, Shaanxi, and Beijing.

Insects likely cause SGS since the insecticide spray significantly reduces SGS incidences [17, 30, 31]. The bean bug *R. pedestris* are susceptible to insecticide treatment [17], but the SGS caused by *R. pedestris* is unrelated to plant virus infection [12]. We showed that *R. pedestris* could not transmit the virus to soybean plants, even though they could carry SoSGV, which is likely taken from the plant sap. In addition to the bean bug, our finding suggested that the level of SoSGV infection should also be sensitive to insecticides because the insecticides can reduce the leafhopper vector population. Thus, maybe challenging, integrated pest management concerning leafhoppers and bean bugs should be implemented to deal with soybean SGS in China.

Leafhoppers are vectors of various plant viruses infecting major crops like rice, maize, wheat, and potato [32]. Among ~ 20,000 species described for leafhoppers, about 50 species were shown to be able to transmit plant viruses [33]. The common brown leafhopper (*O. orientalis*) is the only species we found to survive on soybean in the experimental conditions that are able to transmit SoSGV. The common brown leafhopper can also transmit tobacco yellow dwarf virus (*Mastrevirus, Geminiviridae*) to beans [34] and tobacco [35–38]. Since the SoSGV coat protein shares high homology with mastreviruses [29], and the coat protein determines the insect vector specificity [39], the phylogenetic relations of SoSGV coat protein may explain why leafhopper, other than whitefly [13], transmits SoSGV. Leafhoppers are shown to transmit plant geminiviruses in a persistent circulative manner [40], and the common brown leafhopper has a comparatively broad host plant range [41]. These features likely make the common brown leafhopper an effective insect vector and also a trouble for future soybean SGS management.

Legume plants besides soybean were identified as hosts of SoSGV as their sequences were found in the defective DNA in soybean samples (Fig. 6D). The foreign sequence integration is possibly taken place during viral replication. The same approach was used to find possible insect vectors in this study, but no leafhopper sequence or other currently known insect sequences were found. This result suggests that plant hosts be more important for SoSGV to overwinter and likely play a vital role in the viral transmission cycle. In this respect, a full-scale investigation of the natural hosts of SoSGV is needed for better control of this virus.

## Conclusion

Based on the screen of insects captured from soybean fields showing SGS symptoms and the caged feeding experiments, we demonstrated that the leafhopper is the insect vector that transmits SoSGV to legume plants.

## Materials and Methods

### Field Samples Collection, Processing, and Virus Detection

The soybean samples showing stay-green symptoms were collected from Haidian district of Beijing, Fenyang of Shanxi, Yan’An of Shaanxi, Suzhou in Anhui, and Nanjing in Jiangsu from September to October 2022. The total DNA of the sample was extracted with CTAB (A600108, Sangon Biotech, Shanghai, China) and then purified with FastPure Gel DNA Extraction Mini Kit (DC301, Vazyme, Nanjing, China). PCR was conducted with one or two pairs of specific primers for detecting the existence of SoSGV in each sample. The PCR products were separated with 1% agarose gel. The sample was considered SoSGV-infected when tested positive in PCR assay with at least one pair of primers. The primers used in this study are listed in Supplementary Table 1. The results of each sample are listed in Supplementary Table 2.

### Viral Genome Sequencing

Several samples from each province were randomly selected for genome sequencing. The near-complete full-length genomic DNA was amplified with primer #2914 and #2915 by PCR. The PCR products were inserted into the linearized plasmid pGD-C-Flag [42] with ClonExpress II One Step Cloning Kit (C112, Vazyme, Nanjing, China). Ligation products were transformed into *E. coli* by heat shock method. One colony of each sample was picked for sanger sequencing. To obtain the complete viral genome sequences, another PCR using primers #3133 and #3134 was conducted to amplify the region containing the primers used in the first round of PCR. A total of 22 full-length genomic sequences of SoSGV were obtained. The Genbank accession numbers (OQ079694 - OQ079715) and sequences were listed in Supplementary Table 3. The genomic sequences of defective SoSGV were also obtained with the same method and listed in Supplementary Table 4.

### Sequences Alignment and Phylogenetic Analysis

Six viral genomic sequences (MT221447.1, OM914609.1, OL404964.1, OL855975.1, OM145986.1, and MZ505080.1) were obtained from NCBI. Although they were named differently [27–29], they all belong to the species termed SoSGV in this and the former study [13]. The 6 sequences were aligned with the 22 genomic sequences obtained in this study using MAFFT (version 7.475). The unrooted phylogenetic tree was reconstructed by using the Maximum Likelihood method and Tamura-Nei model [43] with MEGA (version 11.0.11) [44]. All the sites were used in the calculation. The tree with the highest log likelihood (−6848.70) is shown. A discrete Gamma distribution was used to model evolutionary rate differences among sites (2 categories (+G, parameter = 0.1475)). The tree is drawn to scale, with branch lengths measured in the number of substitutions per site. The phylogeny test was conducted using the Bootstrap method with 100 replicates. The sequence identity between the isolates was calculated by Clustal (version 2.1) (https://www.ebi.ac.uk/Tools/msa/clustalo/).

### Insects Capture and Virus Detection

The insects from Anhui province were all caught with an insect net in the soybean field in Suzhou, Anhui, in October 2022. The leafhoppers from Jiangsu were caught in the grass by a soybean field in Nanjing in November 2022. The insects were reared in a nylon cage (30 cm × 30 cm × 30 cm, pore size 0.12 mm).

The total DNA of the insect was extracted from the intact insect with CTAB and purified, as mentioned above in the Materials and Methods section. PCR using primers #3133 and #3134 was used to check the existence of SoSGV. For *R. pedestris* and leafhoppers, the genes coding for Actin and Histon 3 were used as internal loading control, respectively.

### Virus transmission assay

*Glycine max* cv. ‘Nannong 1138-2’, *Glycine soja* varieties ‘Yong-10’, ‘Yong-27’, and ‘Yong-40’, *Vigna unguiculata* cv. ‘Te xuan zhang tang wang’ and a *Vigna angularis* landrace were used in the virus transmission assays. The plants were grown in flowerpots and were put into the nylon cage at the proper stage to carry out the virus transmission experiments. The whole experiments were carried out in the greenhouse with a temperature of 24 ± 2°C, a light period of 14/10 h (day/night), a light intensity of about 10000 lux, and a humidity of 60 ± 10%.

After the insects infested the plants, leaf samples from different individual plants were sampled and treated as biological repeats. The plants grown free of insects were used as mock controls. DNA extraction and virus detection were conducted as former descriptions with primer #3133 and #3144. The water and the plasmid containing SoSGV genomic DNA were used as negative and positive controls, respectively. The genes coding for β-tubulin of each plant species were used as internal loading controls. In the qPCR analysis, ChamQ SYBR qPCR Master Mix (Q341, Vazyme, Nanjing, China) was used with the StepOne™ Real-Time PCR System (Applied Biosystems, Foster City, USA). The relative accumulation of the virus was calculated by a 2^-ΔΔCT^ method [45].

### High Throughput Sequencing and Data Analysis

The plant samples from Beijing, Shanxi, and Anhui were used to amplify the fragments of defective SoSGV with primers #3058 and #3059. PCR products of each sample were mixed and separated in an agarose gel electrophoresis. The 200-600 bp products were recovered using FastPure Gel DNA Extraction Mini Kit (DC301, Vazyme, Nanjing, China) and sequenced by the second-generation Illumina PE300 system. The paired reads were assembled with PEAR (version 0.9.6). The assembled sequences and the remaining unassembled reads were treated with HISAT2 (version 2.2.1) to remove the sequences that matched SoSGV The remaining sequences were clustered by the similarity threshold of 0.9, and one sequence from each cluster was combined to build the unique sequence database with CD-HIT (version 4.8.1). Finally, the online tool BLAST (https://blast.ncbi.nlm.nih.gov/Blast.cgi) was used to identify the possible species to which the unique sequences belong. The sequences harboring any identified insertion sequence were listed in Supplementary Table 5.

## Author contributions

Conceptualization, X.K., L.K. and Y.J.; Methodology, X.K., Y.J.; Investigation,Y.J., X.K., H.Z., X.S., H.X., Q.Y., C.X.; Resources, X.K., L.K., Z.H., S.W., and W.L.; Writing, X.K., and Y.J.; Supervision, X.K. and L.K.; Funding Acquisition, X.K. and L.K. All authors read and approved the final manuscript.

## Acknowledgments

The authors acknowledge Changfa Zhou (Nanjing Normal University) for assisting in identifying insect species, Kai Zhang (Hebei Normal University of Science & Technology) for sharing the *G. soja* varieties, Shi Sun (Chinese Academy of Agricultural Sciences), Junkui Ma (Shanxi Academy of Agricultural Sciences), and Fuqin Liang (Yan’an Institute of Agricultural Sciences) for collecting soybean samples.

## Availability of data and materials

All data are available within the article and its supplementary materials.

## Ethics approval and consent to participate

Not applicable

## Consent for publication

Not Applicable.

---

**Supplementary Figure 1.**
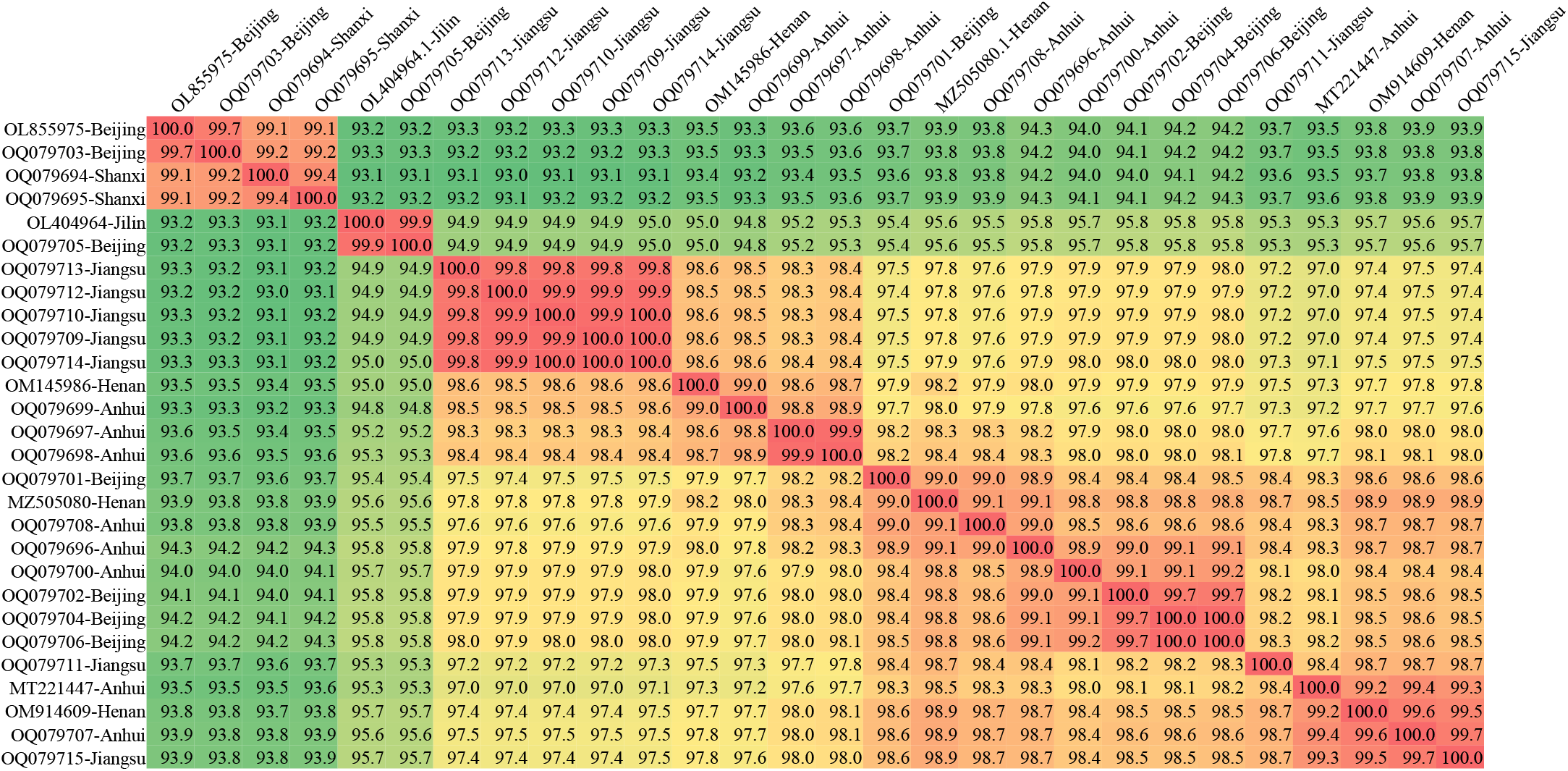
The percent identity matrix of aligned SoSGV genomes. The genome sequences of different SoSGV isolates were used to calculate the identity between each pair of the SoSGV isolates. The calculated percent identity matrix is shown.

**Supplementary Figure 2.**
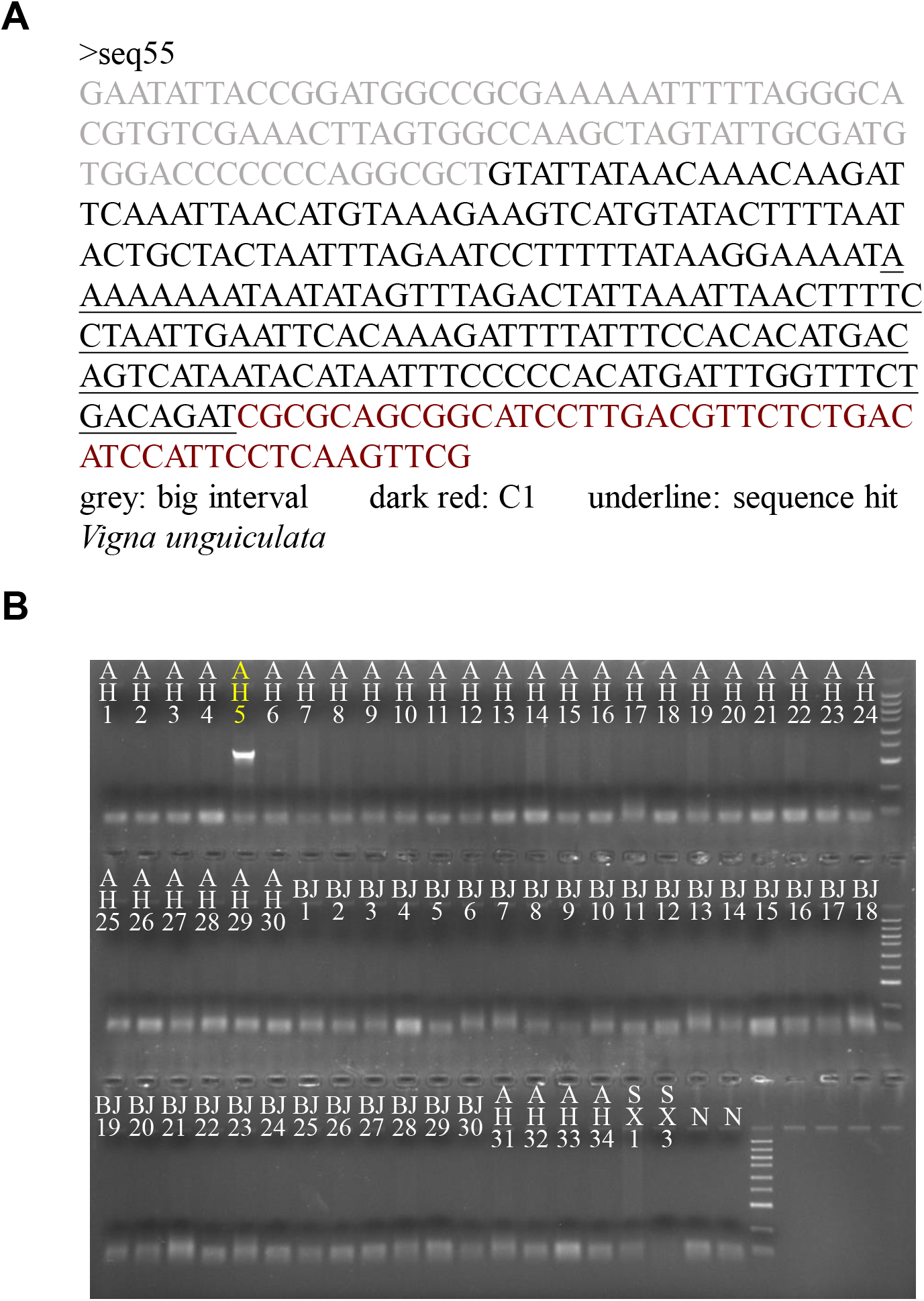
PCR amplification of the SoSGV defective DNA containing sequences from cowpea *Vigna unguiculata*. (A) A fragment of sequence from cowpea inserted in the SoSGV defective DNA clone seq55. (B) PCR amplification of the cowpea sequence from soybean samples showing SGS symptoms using primers #3240 and #2907. Sample #5 collected from Anhui Province (AH5) is detected as positive.

**Supplementary Figure 3.**
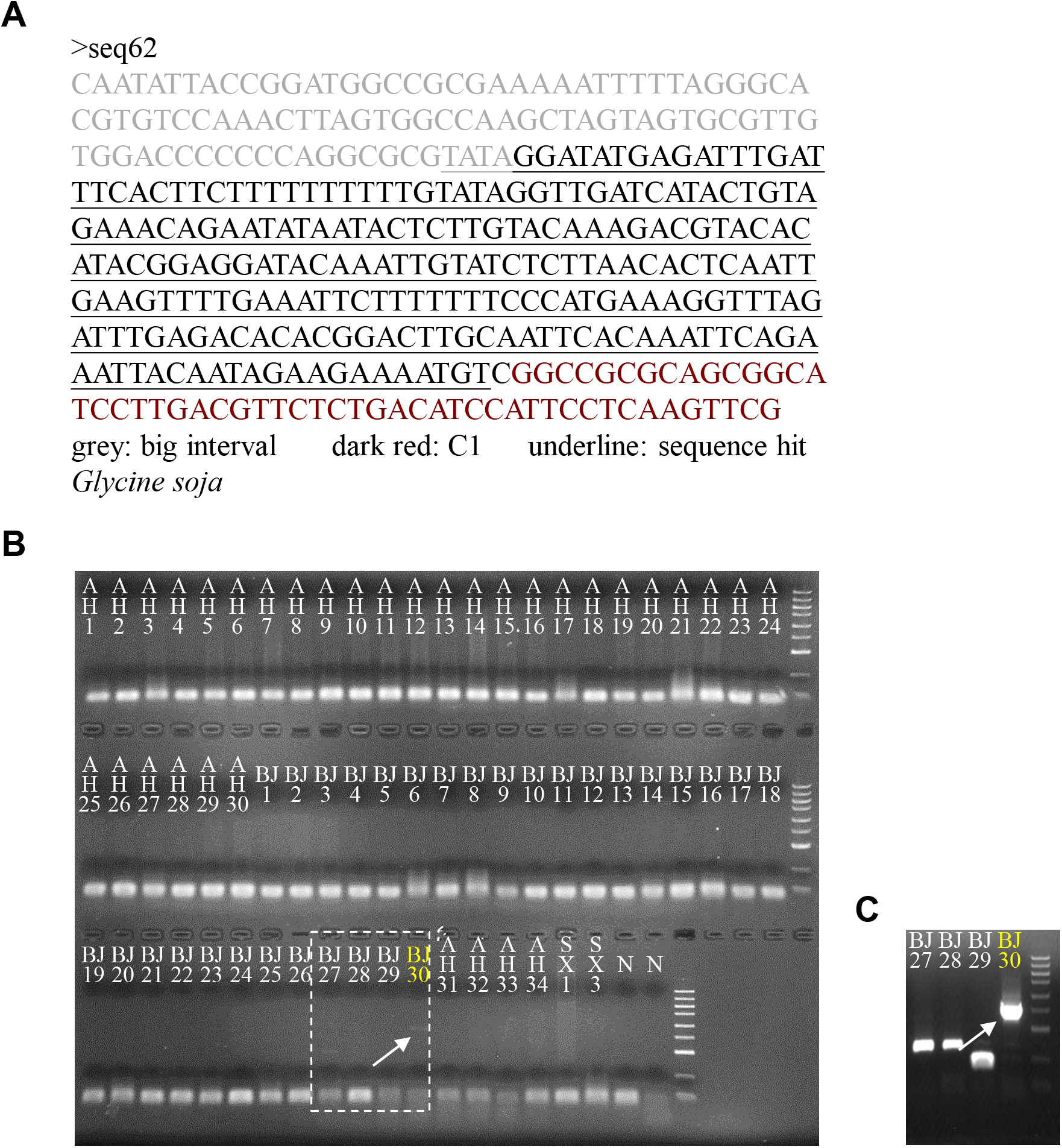
PCR amplification of the SoSGV defective DNA containing the sequence from wild soybean *Glycine soja*. (A) A fragment of sequence from *Glycine soja* inserted in the SoSGV defective DNA clone seq62. (B) PCR amplification of the *Glycine soja* sequence from soybean samples showing SGS symptoms using primers #3241 and #2907. Sample #30 collected from Beijing (BJ30) is detected as weak positive. (C) A nested PCR assay confirmed the presence of the *Glycine soja* sequence in sample BJ30 by using the PCR products #27 – #30 shown in panel B as templates and another primer pair set (#3241 and #3134).

**Supplementary Figure 4.**
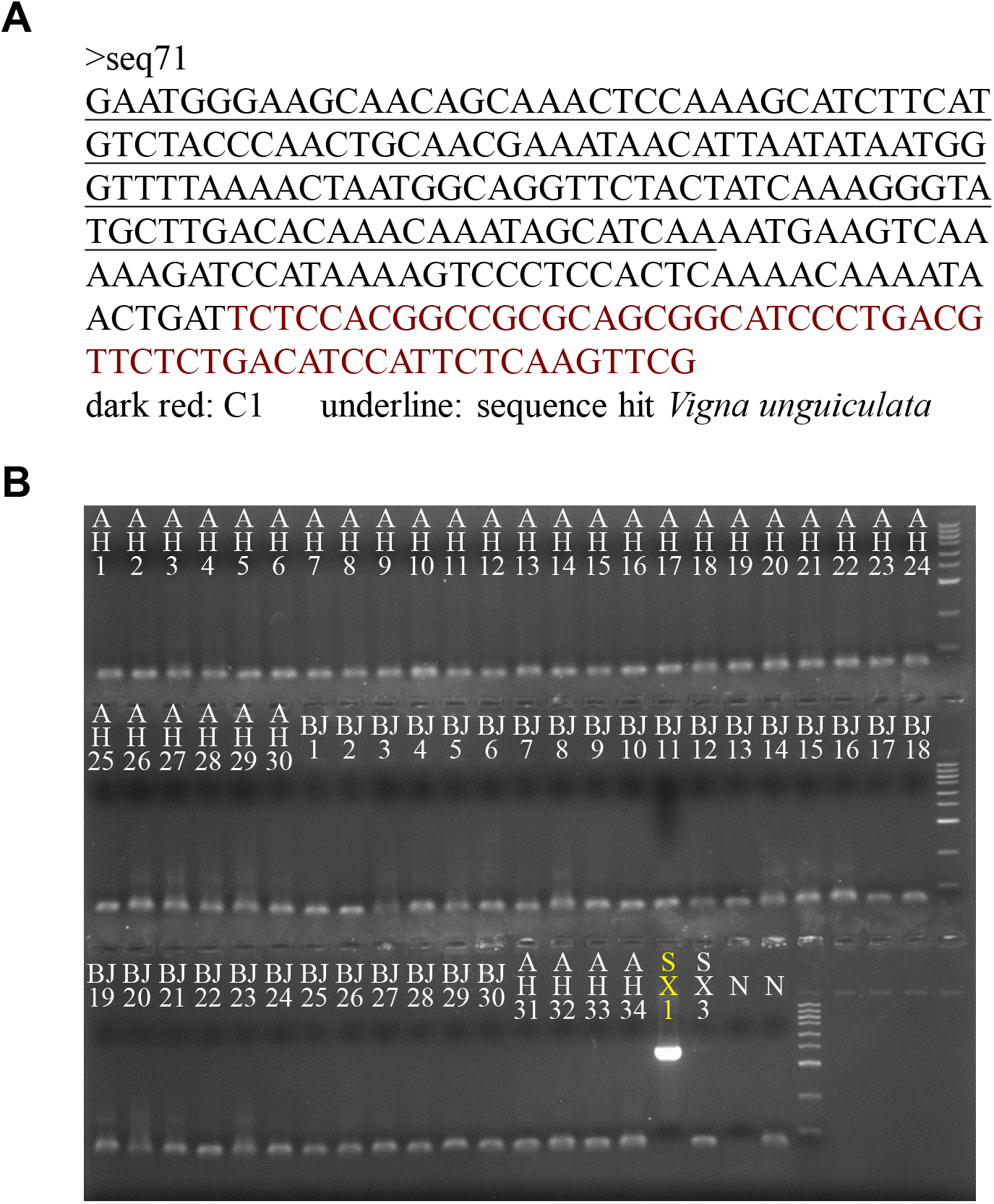
PCR amplification of the SoSGV defective DNA containing the sequence from cowpea *Vigna unguiculata*. (A) A sequence fragment from *Vigna unguiculata* was inserted in the SoSGV defective DNA clone seq71. (B) PCR amplification of the *Vigna unguiculata* sequence from soybean samples showing SGS symptoms using primers #3242 and #2907. Sample #1 collected from Shanxi Province (SX1) is detected as positive.

